# Simulating brain signals with predefined mutual correlations – a technical note

**DOI:** 10.1101/2021.06.01.446620

**Authors:** Alexander Moiseev

## Abstract

**Objective:** When modeling task-related human brain activity it is often necessary to simulate brain signals with specific mutual correlations between them. The signals should resemble those observed in practice, and consist of an “evoked” (“phase-locked”) component and a random oscillatory part. To be neurophysiologically plausible their waveforms must be shaped in a certain way or exhibit specific global features; in technical terms - they should be modulated by a certain envelope function. The goal of this technical note is to describe a simple way of how such signal sets can be obtained.

**Methods:** We derive a procedure which allows generating multi-epoch signals with the above properties. This is done by mixing a “seed” set of waveforms typically reflecting particular qualities of the target brain activity. As an example, the seed set can consist of realizations of colored noise with desired power spectrum, or can be obtained from real brain measurements.

**Results:** The algorithm yields a set of n multi-epoch signals with specified mutual correlations. Evoked parts, oscillatory parts and global envelopes of the signals can be controlled independently in order to obtain desired properties of the generated time courses.

**Conclusion:** The procedure provides versatile sets of mutually correlated signals suitable for modeling task-related brain activity.

**Significance:** In contrast to other methods often relying on complicated computations, the suggested approach is straightforward and easy to apply in everyday practical work, yet yielding realistic “functionally connected” simulated brain signals.

## I. Introduction

In the field of functional brain imaging by means of magnetoencephalography (MEG) or electroencephalography (EEG) one often needs to model brain activity or to validate various techniques by means of computer simulations. Simulating the human brain in general is a vast area of research; an epic overview of the topic with hundreds of references may be found here [1]. A more narrow task is to model waveforms of electromagnetic fields generated by a set of possibly connected brain sources. There is an abundance of publications here too, so we’ll name just a few. First, a general topic of generating signals with desired spectral, statistical or other parameters is developed in detail in radar and communications literature - see for example [2], [3]. Then there are many works devoted to bioelectromagnetic signals specifically, sometimes using quite complicated approaches involving biochemical cell models or using artificial neural networks [4]–[12]. We should also note a “virtual brain” project that aims at providing a versatile software framework for simulating almost any aspect of brain activity [13], [14].

In this short technical note we only focus on a commonly encountered task of simulating sets of multi-epoch brain signals whose time courses have realistic shapes and predefined mutual correlations. Specifically, such signals may include a part which is identical for all epochs and represents an “evoked” or “phase-locked” component of the brain activity, as well as a random part changing from epoch to epoch – so called oscillatory, or “time-locked” component.

For just a single pair of sources the desired correlation may be readily obtained, for example, by using sinusoidal signals with a properly selected phase difference. However such signals hardly resemble brain activity observed in real life. Simulating larger number of sources with complicated mutual inter-dependencies becomes more challenging. At first glance, a *n*-dimensional signal set with any desired covariance matrix ***R*** is easily created by mixing some standardized uncorrelated “seed” set with any *n* × *n* mixing matrix ***X*** such that ***XX***^*T*^ = ***R***, where the superscript “*T*” denotes matrix transposition. Indeed this would be a solution if the target set included only oscillatory components without any other constrains. When one needs to take into account the evoked parts and also to control typical common shapes of generated epochs the procedure becomes significantly more involved.

This technical note describes a simple linear mixing algorithm which allows generating an arbitrary number of signal time courses (epochs) possessing the above features where for every individual epoch the signals are mutually correlated with a given mathematically consistent correlation matrix. Particular qualities of those signals can be controlled by properly choosing an initial “seed” set of waveforms. For example, those can be obtained from real M/EEG measurements or using colored noise with desired spectral characteristics. The generated epochs can also be shaped or tapered with a given envelope function to model typical onsets and fade outs of the brain signals or to control their other common properties. The required mutual correlations are obtained for the ultimate enveloped signals. Suggested procedure thus provides a fast and efficient way to generate realistic correlated brain signals, and at the same time it does not require complicated computations such as those based on the cell models or artificial neural networks.

## II. Methods

### A. Problem formulation

Assume that we need to construct multi-epoch time courses for *n* sources. An actual number of epochs is irrelevant, because the procedure to be designed will work on per epoch basis. Every epoch is a set of *n* real valued signals *p*_*i*_(*t*), *i* = 1, …*n*, specified at *m* discreet time points *t*_*j*_, *j* = 1, …, *m*. We require *m* >> *n* for the time averaging operations over individual epochs to be statistically robust. Taken together signals *p*_*i*_(*t*) can be represented as a *n* × *m* matrix ***p*** with *n* rows (the “source” dimension) and *m* columns (the “time” dimension). Assume also that signals have fixed “evoked” parts *c*_*i*_(*t*) that do not change from epoch to epoch, and random “time-locked” or oscillatory parts *z*_*i*_(*t*) that differ between epochs. Thus our initial representation of the target signal can be written as

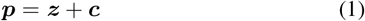

where ***z*** and ***c*** are *n* × *m* matrices.

The following notation will be used further on. Consider an arbitrary set of signals represented by *n* × *m* matrix ***a***. Let source and time domain vectors be *n*-dimensional column and *m*-dimensional row vectors, respectively. Let angle brackets ⟨.⟩ denote time averaging over the course of a single epoch – that is over *m* time points. Then 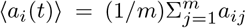. In matrix form we have 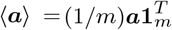, where **1**_*m*_ is a *m*-dimensional row vector of ones. The set ***a*** can be separated into its mean ⟨***a***⟩ and a fluctuating part ***ã***: ***a*** = ⟨***a***⟩ + ***ã***; ***ã*** ≜ ***a*** − ⟨***a***⟩. Finally, let ***C***_*a*_ denote a *n* × *n* covariance matrix of the signal set ***a***, that is ***C***_*a*_ = ⟨***ãã***^*T*^⟩. As matrix multiplication inside the angular brackets already involves summation over the time points, the averaging operation ⟨.⟩ in this case expands as ***C***_*a*_ = ⟨***ãã***^*T*^⟩*/*(*m* − 1) (an unbiased estimate of the covariance). Alternatively 1*/m* (maximum likelihood) normalization can be used; this will not change the final expressions provided the same normalization is applied everywhere. A correlation matrix is a special case of a covariance matrix with all diagonal elements equal to 1. To emphasize that some covariance matrix is in fact a correlation matrix we’ll write ***Ĉ***_*a*_ instead of ***C***_*a*_.

To proceed further let us rewrite (1) in a different form. First without loss of generality we will require that the target signals *p*_*i*_ have a unit variance: 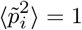; afterwards one can always scale those signals as needed and that will not affect their correlation matrix. Assume also that evoked signals *c*_*i*_ are normalized to have a unit variance: 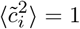. To control relative contributions of phase-locked and time-locked components to the full signal, introduce quotients *q*_*i*_, 0 ≤ *q*_*i*_ ≤ 1, *i* = 1, …, *n* such that 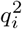 specifies what part of the variance of the full signal 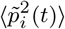 is due to the variance of the evoked part 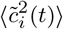. Then instead of (1) each *p*_*i*_(*t*) can now be written as *p*_*i*_(*t*) = *z*_*i*_(*t*) + *q*_*i*_*c*_*i*_(*t*). We want oscillatory parts *z*_*i*_, *i* = 1, …, *n* to be zero mean random signals with specified envelopes, for example we want *z*_*i*_(*t*) to be smoothly tapered at the epoch ends to be physiologically plausible: *z*_*i*_(*t*) = *e*(*t*)*r*_*i*_(*t*), where *r*_*i*_(*t*) is some unconstrained random signal, and *e*(*t*) is the envelope (taper) function: *e*(*t*) = {*e*_*j*_}, *j* = 1, …, *m*. The latter is assumed to be the same for all *n* signals for simplicity. Then in matrix form for the oscillatory part of the signal we have

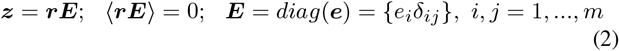

where *δ*_*ij*_ is the Kronecker’s delta.

To get a compact result an additional constrain is needed. Namely we’ll seek a solution where the oscillatory parts of all signals are uncorrelated with all the evoked parts: ⟨***zc***^*T*^⟩ = **0**. Summarizing all of the above we can now write instead of (1):

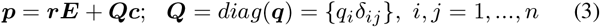

where

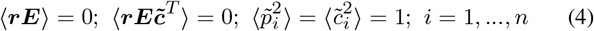

Now our problem can be stated as follows.

**Problem**. Given unit variance evoked signals *c*_*i*_(*t*), quotients *q*_*i*_, 0 ≤ *q*_*i*_ ≤ 1, *i* = 1, …, *n* and an envelope function *e*(*t*), construct a set of random signals ***r***(*t*) = {*r*_*ij*_}, *i* = 1, …, *n, j* = 1, …, *m* such that simulated “brain signals” ***p*** in (3), (4) have desired correlation matrix ***Ĉ***_*p*_.

### B. Solution

Start with an arbitrary (*n* × *m*) *seed signal set* ***s*** which should usually reflect desired features of the modeled brain activity. As already mentioned it can be obtained in different ways, for example from real measurements or by using band-pass filtered or colored noise with desired power spectrum.

Then for every epoch of the seed signal set ***s*** corresponding epoch of the unconstrained signal ***r*** is obtained by the formula:

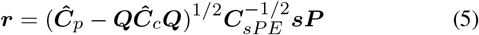

Accordingly an epoch of the full target brain signal ***p*** is given by (3). In (5) ***Ĉ***_*c*_ is the correlation matrix of the evoked parts of the target signal, 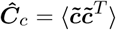; ***P*** is a time domain (*m* × *m*) projection matrix given by the expression:

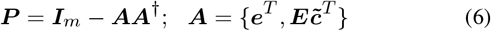

where ***I***_*m*_ is a *m*-dimensional identity operator; *m* × (*n* + 1) rectangular matrix ***A*** is obtained by concatenating ***e***^*T*^ and columns of the product 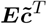; symbol “†” denotes matrix pseudo-inverse. Finally, ***C***_*sPE*_ denotes covariance of the projected and tapered seed set ***s***:

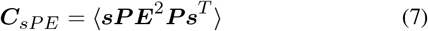

***Remark***. *With straightforward modifications, solution* (5) *–* (7) *can be generalized to the case of complex-valued signals*.

Note that existence of the evoked parts *c*_*i*_(*t*) limits possible choice of the target correlation matrices ***Ĉ***_*p*_, because solution (5) only exists if ***Ĉ***_*p*_ − ***QĈ***_*c*_***Q*** is a positively defined matrix. Indeed, correlations between the evoked signals *c*_*i*_, which are reflected by the term ***QĈ***_*c*_***Q***, impose a lower limit on correlations between the full signals *p*_*i*_(*t*). For example, there always exists large enough ***Q*** such that it becomes impossible to create a completely uncorrelated set ***p*** (i.e. ***Ĉ***_*p*_ = ***I***_*n*_) if evoked parts are at least partially correlated.

A step by step computational procedure implementing solution (5) is outlined in Listing 1. Mathematical proofs are given in *Appendix I*

#### Listing 1.

Computation steps

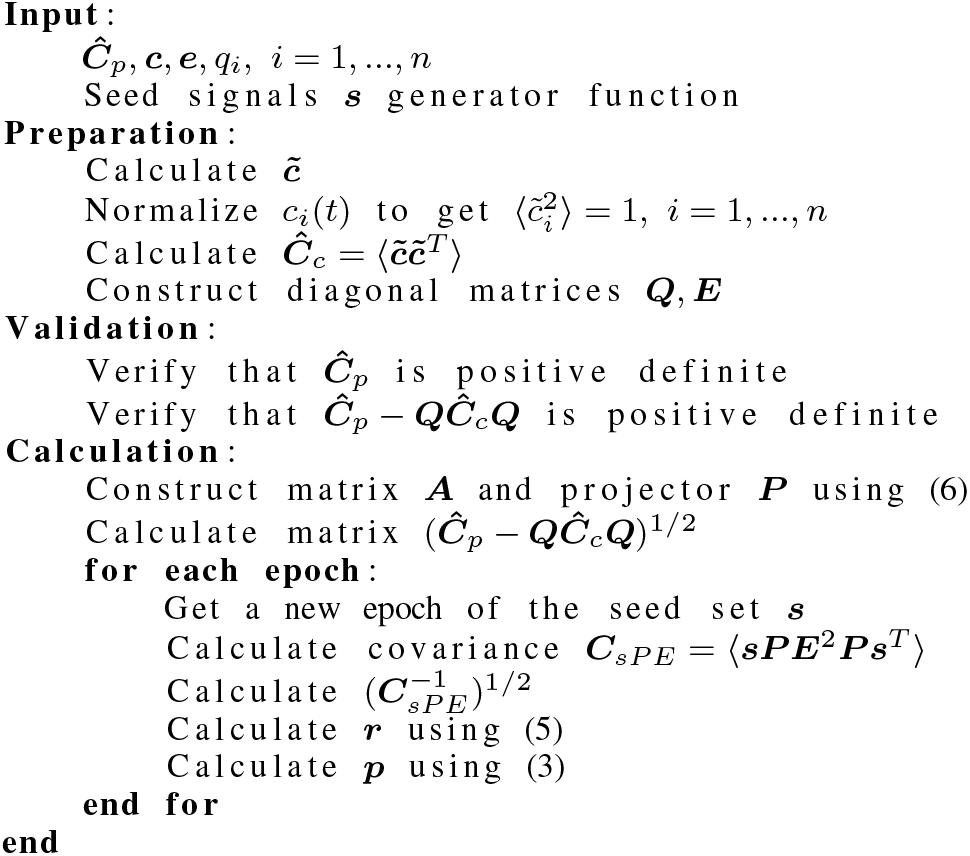

## III. Example

Imagine a situation where we want to model an event-related response from three pairs of functionally related brain locations in three main frequency bands. Let sources {1, 2} represent alpha band activity in 8 to 12 Hz range, sources {3, 4} represent beta activity in 16 to 20 Hz range, and sources {5, 6} model gamma activity in 35 to 40 Hz range.

Assume also that alpha and beta signals carry evoked components, while gamma signals don’t. The evoked components do not need to belong to corresponding frequency bands. For simplicity let the evoked parts of the alpha pair {1, 2} have the same waveforms for both sources but opposite signs, while for the beta pair {3, 4} they are also identical and have the same signs. We want the evoked parts to contribute 60% to the variance of the total generated signals: 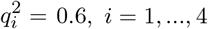.

We would like the full (that is, the oscillatory and evoked parts combined) signals to be correlated, to reflect functional relations between corresponding brain areas. Specifically, we request the following correlation coefficients between the signals of each pair: − 0.9 for alpha, 0.5 for beta and 0.2 for gamma. Further assume that across the pairs alpha and beta signals are uncorrelated, and so are the beta and gamma signals. However we want a moderate correlation of 0.33 within odd-numbered signals and within even-numbered signals of the alpha and gamma pairs - that is between signals 1, 5 and signals 2, 6. The above settings result in the target correlation matrix ***Ĉ***_*p*_ shown in (8)

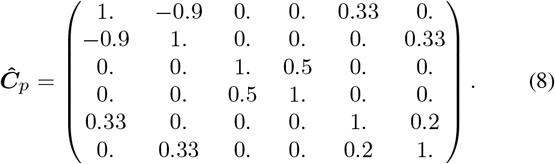

Finally, we use an envelope *e*(*t*) = *h*(*t, T*)*e*^−6*t/T*^ where *h*(*t, T*) is a Hanning window function and *T* is a signal duration; set *T* = 1 *s* for this example. The exponential multiplier is added to account for fading out of the event-related signals with time.

Instances of signals comprising one epoch of the seed, evoked and target sets which were generated in accordance with the above requirements are depicted in Fig. 1; the Python code for this example can be found at https://github.com/aa-moiseev/pub-corr-src-set.git.

**Fig. 1.**
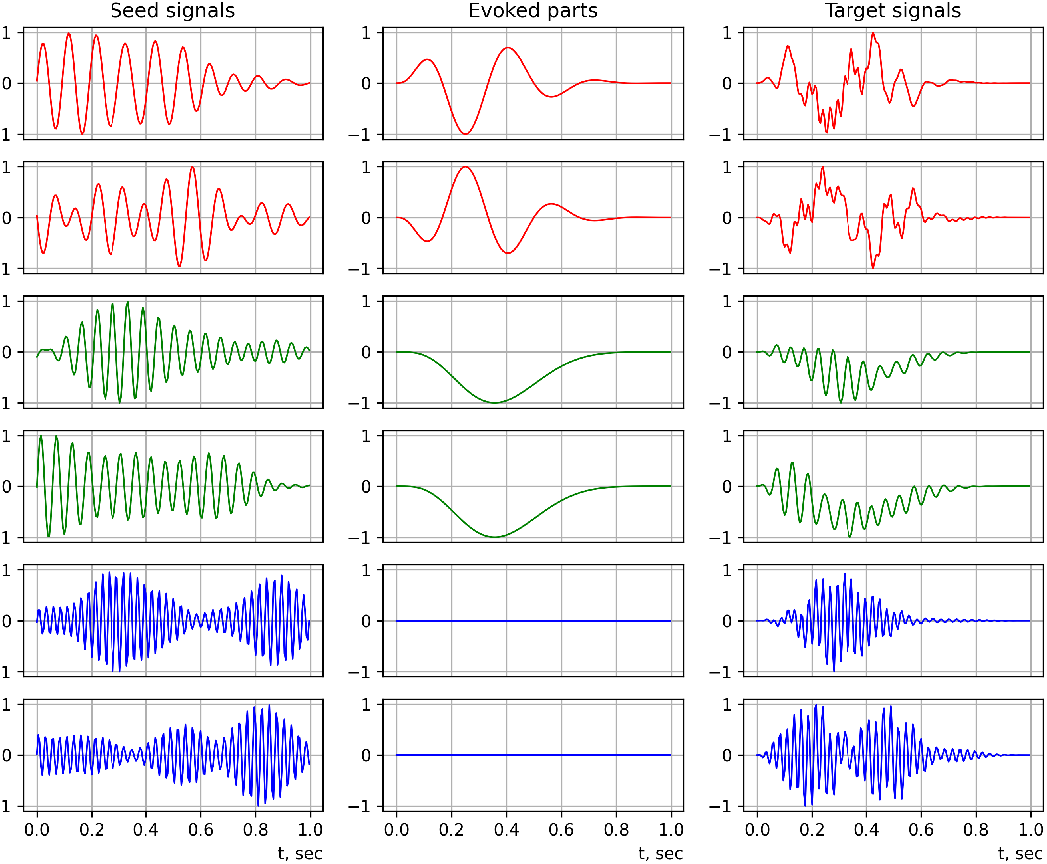
Seed (*s*), evoked (*c*) and target (*p*) signals for alpha (red), beta (green) and gamma (blue) pairs, scaled to [−1, 1] range. The seed signals ***s***_***i***_, ***i* = 1, …6** were produced by passing white noise sampled at **200 *s***^**−1**^ rate through 8-th order Butterworth filter tuned to the corresponding frequency band. The evoked parts were constructed using expressions ***c***_**1**,**2**_ **= ±*e*(*t*) sin(6π*t/T*)** for the alpha pair and ***c***_**3**,**4**_ **= −*e*(*t*)*h*(*t, T*)** for the beta pair.

## IV. Discussion

The resulting set ***p*** shown in the third column in Fig. 1 has a correlation matrix exactly matching the target (8). However, one can notice that due to linear mixing of the seed signals some leakage of gamma frequencies into the alpha band pair *p*_1_, *p*_2_, and of alpha-frequencies into the gamma pair *p*_5_, *p*_6_ occurred. Such leakage cannot be avoided with this type of algorithm if one wants the signals from different bands to be partially correlated. This should be kept in mind when using other, more complicated seed signals for modeling: one may expect specific distinct features of those to be mixed in the resulting set in accordance with dependencies between the signals imposed by ***Ĉ***_*p*_.

Also as explained in *section II-B* a solution wouldn’t exist if matrix (8) was not positively defined. In practice verifying this is important because an arbitrarily compiled ***Ĉ***_*p*_ does not need to be and likely is not positively defined, and therefore cannot represent any realizable set of mutual correlations. For example in our case this happens if a stronger correlation between alpha and gamma pairs is requested – say 0.5 instead of 0.33. An intuitive explanation of this mathematical fact is that for high enough correlation between the alpha and gamma pairs the strong (anti) correlation among the alpha signals ((***Ĉ***_*p*_)_1,2_ = − 0.9) would induce a stronger correlation among the gamma signals, which contradicts the requirement for the gamma pair to be weakly correlated ((***Ĉ***_*p*_)_5,6_ = 0.2).

## V. Conclusion

We developed a simple procedure for generating a set of mutually correlated signals suitable for modeling task-related brain activity. The procedure allows to independently control oscillatory and evoked parts of the signals as well as overall signal envelopes, thus providing a way to create versatile realistic signal sets for brain modeling. The algorithm relies on straightforward linear algebra calculations and can be readily implemented without much programming effort. An example Python code is also provided.

## Appendix I

### Mathematical proofs

Start with verifying that signals (3) satisfy constraints (4). First recall that 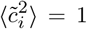 is just a normalization assumption which can always be met by proper scaling. Next check that oscillatory parts have zero means: ⟨***rE***⟩ = 0. Indeed, according to (5) 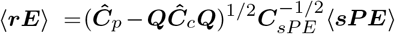, but 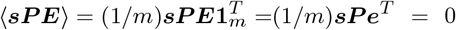, because by construction of the projector ***P*** in (6) ***Pe***^*T*^ = 0. Similarly oscillatory parts ***rE*** are uncorrelated with evoked signals 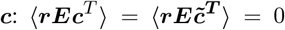, because again by construction of ***P*** we have 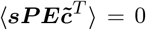. Thus the first two conditions in (4) are verified. Additionally for the varying part of the evoked component of the total signal 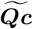 we have 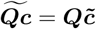, because ***Q*** does not depend on time. It follows then that the oscillatory part of ***p*** is uncorrelated with its evoked part: 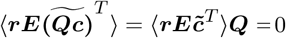.

The third constraint 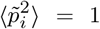 will be satisfied automatically because ***Ĉ***_*p*_ is a correlation matrix. Let us verify that ***Ĉ***_*p*_ is in fact a covariance of the signal ***p***. By definition, covariance of ***p*** is 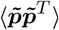. We have

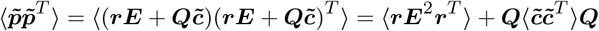

because as we just showed cross-correlations between evoked and oscillatory parts disappear. Then using expression (5) for ***r*** and (7)for ***C***_*sPE*_, we get

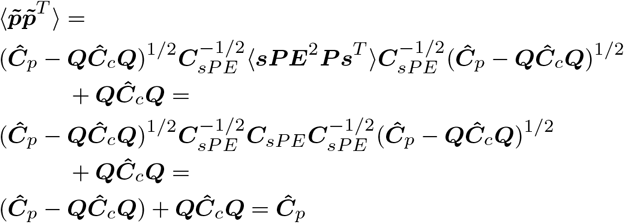

– end of proof.

## References

[1] X. Fan and H. Markram, “A Brief History of Simulation Neuroscience,” Frontiers in Neuroinformatics, vol. 13, p. 32, 5 2019.

[2] M. Viswanathan, Wireless Communication Systems in Matlab, 2nd ed. Independently published (June 2020), 2020.

[3] M. C. Jeruchim, P. Balaban, and K. S. Shanmugan, Simulation of Communication Systems: Modeling, Methodology and Techniques (Information Technology: Transmission, Processing and Storage), 2nd ed. Kluwer academic/plenum publishers, 2000.

[4] A. Anzolin, J. Toppi, M. Petti, F. Cincotti, and L. Astolfi, “SEED-G: Simulated EEG Data Generator for Testing Connectivity Algorithms,” Sensors, vol. 21, no. 11, p. 3632, 2021.

[5] S. A. Neymotin, D. S. Daniels, B. Caldwell, R. A. McDougal, N. T. Carnevale, M. Jas, C. I. Moore, M. L. Hines, M. Hämäläinen, and S. R. Jones, “Human Neocortical Neurosolver (HNN), a new software tool for interpreting the cellular and network origin of human MEG/EEG data,” eLife, vol. 9, p. e51214, jan 2020. [Online]. Available: https://doi.org/10.7554/eLife.51214

[6] S. Haufe and A. Ewald, “A simulation framework for benchmarking EEG-based brain connectivity estimation methodologies,” Brain Topogr, vol. 32, p. 625–642, 2019.

[7] L. R. Krol, J. Pawlitzki, F. Lotte, K. Gramann, and T. O. Zander, “SEREEGA: Simulating event-related EEG activity,” Journal of Neuroscience Methods, vol. 309, pp. 13–24, 2018. [Online]. Available: https://www.sciencedirect.com/science/article/pii/S0165027018302395

[8] E. Hagen, S. Næss, T. V. Ness, and G. T. Einevoll, “Multimodal modeling of neural network activity: Computing LFP, ECoG, EEG, and MEG signals with LFPy 2.0,” Frontiers in Neuroinformatics, vol. 12, 2018. [Online]. Available: https://www.frontiersin.org/article/10.3389/fninf.2018.00092

[9] S. M. Plis, J. S. George, S. C. Jun, J. Paré-Blagoev, D. M. Ranken, C. C. Wood, and D. M. Schmidt, “Modeling spatiotemporal covariance for magnetoencephalography or electroencephalography source analysis,” Phys. Rev. E, vol. 75, p. 011928, Jan 2007. [Online]. Available: https://link.aps.org/doi/10.1103/PhysRevE.75.011928

[10] N. M. Tomasevic, A. M. Neskovic, and N. J. Neskovic, “Artificial neural network based approach to EEG signal simulation,” Int J Neural Syst., vol. 22, p. 1250008, 2012.

[11] N. M. Tomasevic, A. M. Neskovic, and N. J. Neskovic, “Correlated EEG signals simulation based on artificial neural networks,” International Journal of Neural Systems, vol. 27, p. 1750008, 2017.

[12] A. T. Herdman, “SimMEEG software for simulating event-related MEG and EEG data with underlying functional connectivity,” Journal of Neuroscience Methods, vol. 350, p. 109017, 2021. [Online]. Available: https://www.sciencedirect.com/science/article/pii/S0165027020304404

[13] P. Sanz Leon, S. Knock, M. Woodman, L. Domide, J. Mersmann McIntosh, and V. Jirsa, “The Virtual Brain: a simulator of primate brain network dynamics,” Frontiers in Neuroinformatics, vol. 7, 2013. [Online]. Available: https://www.frontiersin.org/article/10.3389/fninf.2013.00010

[14] M. Schirner, L. Domide, D. Perdikis, P. Triebkorn, L. Stefanovski, R. Pai, P. Prodan, B. Valean, J. Palmer, C. Langford, A. Blickensdörfer, M. van der Vlag, S. Diaz-Pier, A. Peyser, W. Klijn, D. Pleiter, A. Nahm, O. Schmid, M. Woodman, L. Zehl, J. Fousek, S. Petkoski, L. Kusch, M. Hashemi, D. Marinazzo, J.-F. Mangin, A. Flöel, S. Akintoye, B. C. Stahl, M. Cepic, E. Johnson, G. Deco, A. R. McIntosh, C. C. Hilgetag, M. Morgan, B. Schuller, A. Upton, C. McMurtrie, T. Dickscheid, J. G. Bjaalie, K. Amunts, J. Mersmann, V. Jirsa, and P. Ritter, “Brain simulation as a cloud service: The Virtual Brain on ebrains,” NeuroImage, vol. 251, p. 118973, 2022. [Online]. Available: https://www.sciencedirect.com/science/article/pii/S1053811922001021

